# RNA Pore Translocation with Static and Periodic Forces: Effect of Secondary and Tertiary Elements on Process Activation and Duration

**DOI:** 10.1101/2021.01.26.428218

**Authors:** Matteo Becchi, Pietro Chiarantoni, Antonio Suma, Cristian Micheletti

## Abstract

We use MD simulations to study the pore translocation properties of a pseudoknotted viral RNA. We consider the 71-nucleotide long xrRNA from Zika virus and establish how it responds when driven through a narrow pore by static or periodic forces applied to either one of the two termini. Unlike the case of fluctuating homopolymers, the onset of translocation is significantly delayed with respect to the application of static driving forces. Because of the peculiar xrRNA architecture, activation times can differ by orders of magnitude at the two ends. Instead, translocation duration is much smaller than activation times and occurs on timescales comparable at the two ends. Periodic forces amplify significantly the differences at the two ends, both for activation times and translocation duration. Finally, we use a waiting-times analysis to examine the systematic slowing-downs in xrRNA translocations and associate them to the hindrance of specific secondary and tertiary elements of xrRNA. The findings ought to be useful as a reference to interpret and design future theoretical and experimental studies of RNA translocation.

## Introduction

RNA translocation is a rapidly growing avenue in theoretical and experimental single-molecule studies. Pore translation has recently allowed for distinguishing different types of tRNAs[1] and quantifying mRNA expression[2]. Measurements of ionic current blockades in nanopores[3] have been used to sequence RNAs[4], to probe salient features of their folding pathway[5] and to detect modified nucleobases[6]. Molecular dynamics simulations have shown that driving RNAs through pores of appropriate width can relay information about their compliance to structural deformations[7] and directional mechanical resistance[8].

RNA translocation properties are of direct biological relevance, too, as they determine the interaction with and response to precessive enzymes. For instance, sequences and structures of viral RNAs have evolved to introduce specific ribosomal slippages needed to produce alternative transcripts [9–12]. Arguably, the most striking example of viral RNA hindrance to enzymatic translocation is given by xrRNAs. These molecules are about 70 nucleotides long, rich in pseudoknots and can resist degradation by exonucleases while remaining processable by other enzymes[13–18].

In a recent theoretical and computational study from our group [8], we used atomistic simulations of Zika xrRNA translocation to explore the microscopic origin of its resistance to degradation. The xrRNA structure, see Fig. 1, is organised so to produce very different structural deformations when one or the other termini are engaged and translocated through the pore. These directional dependent deformations allow the xrRNA to withstand translocation very differently at the two ends. By far, the largest hindrance is encountered at the 5 terminal, which is the same one attacked by exonucleases[15]. Such major directional response, is arguably what makes xrRNA resistant to degradation (initiating at the 5 end) while still processable for transcription and replication (initiating at the 3 end). For both its biological relevance and atypical density of pseudoknots, xrRNA is an ideal substrate to study how the translocation process depends on intrinsic properties, such as secondary and tertiary elements, and extrinsic ones, such as the use of static or periodic pulling modes.

**FIG 1.**
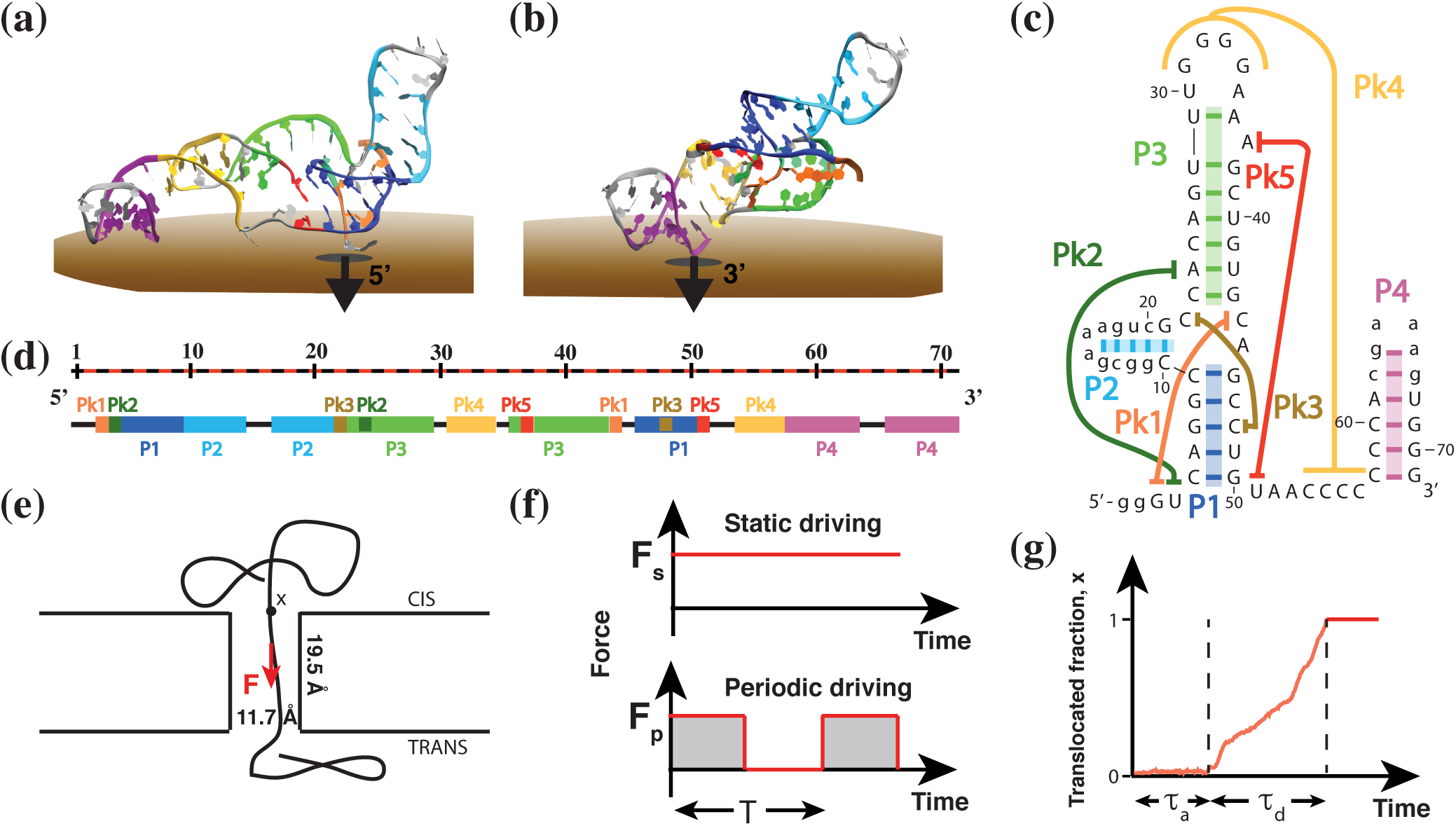
System setup. Typical configurations at the beginning of translocation simulations from the (a) 5^*′*^ and (b) 3^*′*^ ends of Zika xrRNA. The molecule is organised in four helices (P1-P4) and five pseudoknots (Pk1-Pk5), as represented in panels (c) and using the same color-code of panel (a). Specifically, in the two-dimensional graph of panel (c), the backbone connectivity is subsumed by the sequential numbering of the nucleotides, which are indicated with their one-letter code, and the colored dashes and lines indicate the main pairings and interactions of helices and pseudknots. In panel (d), the same motifs are annotated along the one-dimensional representation of the xrRNA. During translocation, the molecule is driven through a cylindrical pore by means of a static or periodic force (f) that is distributed and applied on the *P* atoms that are inside the pore. (g) The translocation progress is monitored via the fraction of translocated atoms, *x*, whose time evolution is sketched in panel (g) along with the indication of the activation time, *τ*_*a*_ and duration, *τ*_*d*_ of the translocation process. The scheme in panel (c) is adapted with permission from ref. 8.

Studying the role of secondary and tertiary elements is important from the polymer physics point of view, as it aptly complements the now well-established theory of translocating homopolymers [19–21]. The latter enjoy a large conformational freedom, and their out-of-equilibrium translocation response can significantly depend on how tension propagates along the fluctuating backbone. By contrast, folded RNAs are structurally constrained by intra-molecular interactions, including base pairings, that introduce translocation barriers that have no counterpart in homopolymers.

The effect of using different driving modes is of interest, too, for at least two reasons. First, to our knowledge, it has not been considered before in connection with RNAs. Second, periodic driving offers a simplified model of translocation as operated by precessive enzymes, which pull on the substrate intermittently.

Here we will address these largely unexplored avenues using molecular dynamics simulations on a native-centric atomistic model of Zika xrRNA. In our study, which adopts the same setup of ref.8, we study how the xrRNA responds when driven from either of its ends through a narrow pore by static and periodic forces. In particular we examine the translocation duration and activation times, and discuss how they significantly vary with the magnitude and period of the driving force, and with pulling directionality, i.e. whether one or the other xrRNA ends are initially engaged. Finally, we examine the so-called waiting times profiles to rationalise the systematic non-uniformities of the translocation process and relate them to the hindrance offered by xrRNA’s secondary and tertiary motifs.

## METHODS

### xrRNA structure

We considered the 71-nucleotide long Zika xrRNA structure of PDB[22] entry 5TPY [15], which is shown in Fig. 1. The molecule features five relatively short pseudoknots, labelled Pk1 to Pk5, mostly concentrated at the 5′ end, and four helices, P1 to P4, see Fig. 1. Two- and one-dimensional representations of the secondary and tertiary motifs are given in Fig. 1c,d.

### System setup

Following ref. 8, the 5′ and the 3′ xrRNA ends were separately primed at the entrance of a narrow cylindrical pore embedded in a parallelepiped slab, see Fig. 1a,b,e. The pore is 11.7 Å wide and 19.5 Å long, approximating the lumen of the Xrn1 exoribonucleases[23].

To treat xrRNA intra-molecular interactions, we used SMOG [24, 25], an implicit-solvent atomistic force field that is native centric. As such, the potential energy, which includes bonded and non-bonded interactions, angular and dihedral terms, are designed to stabilize the native conformation. Excluded volume interactions of the xrRNA with the pore and slab walls were accounted for with truncated and shifted Lennard-Jones potentials.

Constant temperature (Langevin) translocation simulations were carried out with the LAMMPS package [26] after conversion of the input and topology files using the “SMOG-converter” that is publicly available on the github repository[27]. Following ref. 8 the system temperature and energy scale were calibrated by matching the typical stretching forces (~ 15 pN) required to unfold small RNA helices at 300 K. Simulations were carried out with proper atomic masses and with default values of the friction coefficient.

The characteristic simulation time is 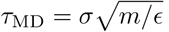 where the typical range and strength of interaction potentials are *σ* = 4 Å and *ϵ* = 0.1 Kcal/mol, respectively, and *m* = 15 amu [8]. The integration time step was set equal to 8.7 · 10^*−*4^ *τ*_MD_. The nominal mapping of simulation units to real units yields *τ*_MD_ ~1.3 ps, though the absence of explicit solvent interactions is expected to skew the model dynamics to being faster than it actually is by orders of magnitude [28]. For this reason, temporal durations are expressed in units of *τ*_MD_ throughout the study.

After a preliminary relaxation run, the molecule was translocated by an external force. We considered two different driving protocols: one with a static force, *F*_*s*_, and one with periodic force switched regularly between 0 and *F*_*p*_ (square-wave). To mimic electrokinetic translocations, the driving force was applied only to the *P* atoms inside the pore. Because the latter can fluctuate in number, the total force, *F*_*s*_ or *F*_*p*_, was equally subdivided among the *P* atoms in the pore. For each considered value of the force and switching rate, we collected from 20 to 40 independent runs.

### Observables

Progress of the translocation process was monitored via the translocated chain fraction, *x*, defined as the fraction of xrRNA atoms that have left the *cis* region, and thus are either in the pore or in the *trans* region.

The translocation activation time, *τ*_*a*_ = *t*_start_ −*t*_0_, measures the time elapsed from the start of the simulation, *t*_0_, to when the leading P atom reaches the *trans* region without retracting from it, *t*_start_. This condition corresponded to the onset of irreversible translocations for all combinations of static and periodic drivings. The translocation duration, *τ*_*d*_ = *t*_end_−*t*_start_, measures the time elapsed from the process activation, *t*_start_ to when the last xrRNA atom enters the *trans* region, *t*_end_.

To characterize the translocation hindrance of different xrRNA regions, we measured the so-called waiting time [29] for each nucleotide. This observable, *w*, is the cumulative time that a nucleotide spends straddling the pore entrance, i.e. with part of its atoms in the *cis* region and others inside the pore. Such “straddling time” may be cumulated over multiple time intervals (in case of retractions) and is averaged over different simulations.

## RESULTS AND DISCUSSION

Recent work from our group has demonstrated a strong directional response of Zika xrRNA to translocation, with the 5′ end offering much more resistance than the 3′ one. The enhanced resistance originates from the peculiar geometry of the pseudoknotted 5′ end, which is encircled by secondary elements that tighten up when the driving force pulls them against the pore rim[8]. These results were established using a force-ramping protocol, a common setup in force-spectroscopy experiments.

Here, we consider different pulling protocols, first a static mode with a constant driving force, *F*_*s*_, and then a periodic mode, where the driving force is switched between 0 and *F*_*p*_ at regular intervals of duration *T/*2, see Fig. 1f.

### Static driving

#### Activation time

For general homopolymers, translocation initiates as soon as the driving force is applied, provided that the latter overcomes the chain’s entropic recoil. This is not the case for the considered system, where intramolecular interactions, such as base pairings, allow the molecule to withstand the exerted force and delay the onset, or activation, of translocation.

This is illustrated by the typical translocation curves of Fig. 2, portraying the time evolution of the translocated chain fraction, *x*, for a constant force *F*_*s*_ = 210 pN.

**FIG 2.**
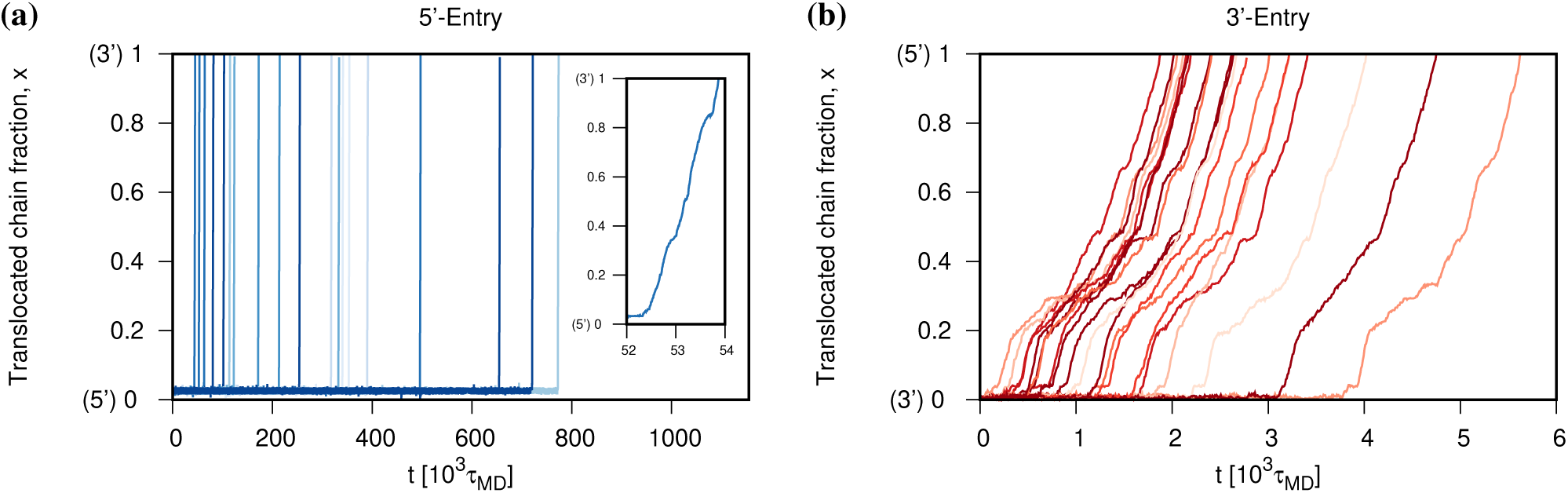
Translocation curves at constant driving. Translocation curves at constant force, *F*_*s*_ = 210 pN, for 5′ (blue) and 3′ (red) entries of Zika xrRNA. 20 independent trajectories are shown in both cases. Because the activation time at the 5′ end is much longer than the duration of translocation, the post-activation *x*(*t*) curves are too steep to show discernible features. The latter can be appreciated in the inset, which shows the post-activation *x*(*t*) curve for a single run.

The *x*-axis scales of the two graphs reveals a striking difference of activation times, *τ*_*a*_, at the two xrRNA ends: the average *τ*_*a*_ is 1.25 10^3^ *τ*_MD_ for 3′-entry and 294 10^3^ *τ*_MD_ for 5′end, a two order-of-magnitude difference. Note that the translocation process at the 3′ end is not only initiated, but also completed in a timespan much smaller than the activation time at the 5 end. This implies that the entire molecule, including the portion resisting translocation at the 5′ end, is disrupted significantly faster from the 3′ end.

A systematic comparison of the activation times, *τ*_*a*_, for 5′ and 3′ pore entries is given in Fig. 3. The linear trends in the semi-log plot of Fig. 3a indicate that *τ*_*a*_ decays about exponentially with the applied force at both ends, 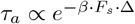. Thus, notwithstanding the complex interactions of the xrRNA termini and the pore, the activation of translocation can be modelled as a two-state process involving a free-energy barrier of effective widthΔ.

**FIG 3.**
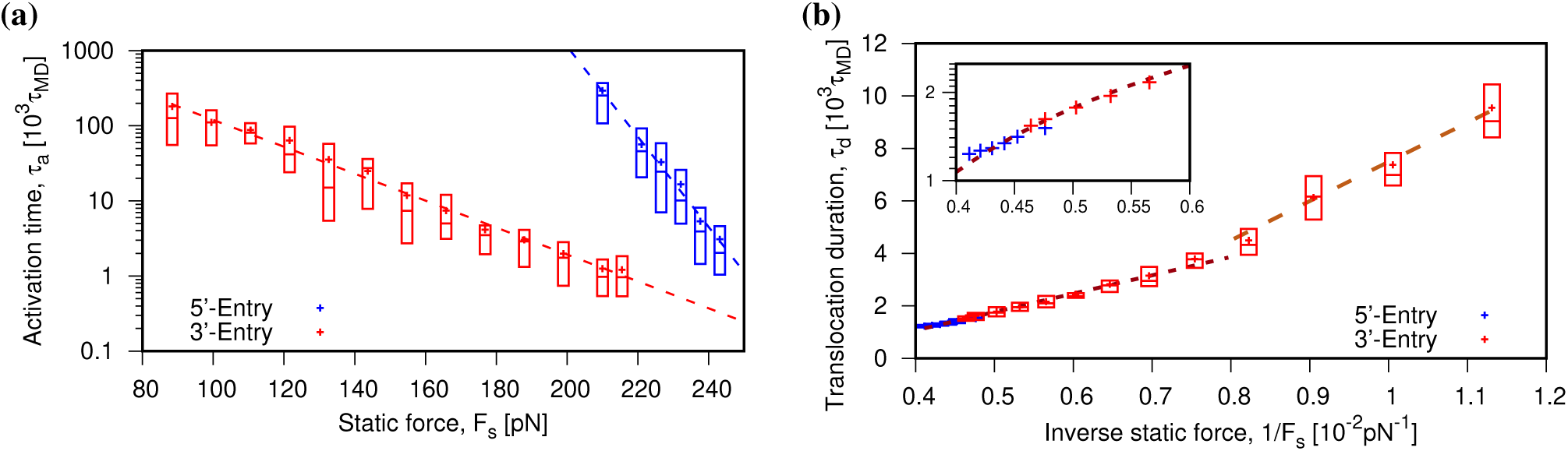
Translocation activation times and process duration at constant driving. Box plots for (a) translocation activation times, *τ*_*a*_, and (b) translocation duration, *τ*_*d*_ for 5′ (blue) and 3′ (red) entries for various static driving forces, *F*_*s*_ (dot, average; center line, median; box limit, upper and lower quartile). The dashed lines in (a) are exponential best fits for *τ*_*a*_ and correspond to barrier widths ofΔ_5′_ = 5.6 0.2 Å andΔ_3′_ = 1.69 0.04 Å. Panel (b) illustrates the dependence of *τ*_*d*_ on the inverse force. Two main regimes are apparent for the 3^*1*^ case; their linear best fits (dashed lines) yield a crossoverat 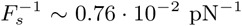 The response at small 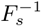 values, i.e. large forces, is highlighted in the inset, where the distinct trends of the 5′ and 3′ entries is better appreciated.

The exponential best fits of the *τ*_*a*_ data, shown by the dashed lines in Fig. 3a, yieldΔ_5′_ = 5.6± 0.2 Å for the 5′-entry andΔ_3′_ = 1.69 ± 0.04 Å for the 3′ one. The values of the effective barrier widths are similar to those established from the Bell-Evans analysis of force-ramped translocations [8], 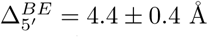 and 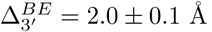

The larger value ofΔ for 5′ entries accounts for the fact that 5′ activation times become progressively larger than 3′ ones as *F*_*s*_ is lowered. The *τ*_*a*_ difference grows to several orders of manitude when *F*_*s*_ is extrapolated to 50-100 pN. Such forces are comparable to those exerted by the most powerful molecular motors[30], and thus the very different activation times are consistent with the peculiar resistance of xrRNA to degrading enzymes, which engage the 5′ end, while the same molecule can be processed from the 3′ end by replicases and reverse-transcriptates.

#### Translocation duration

We now focus on how translocation progresses once initiated. Again, it is informative to contrast the observed behaviour with that of standard homopolymers, where the typical time required to translocate a fraction *x* of the chain scales asymptotically as 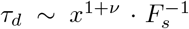 [19, 20, 31-34], where *ν* is the metric exponent. The curves of Fig. 2 depart qualitatively from this scaling law, because the translocation does not proceed uniformly, but presents systematic pauses and slowing downs in correspondence of precise xrRNA regions. We will discuss in more depth these properties further below where we connect them to the secondary and tertiary xrRNA organization. Here, we instead consider the overall duration of the translocation process, *τ*_*d*_, and its dependence on the applied force, *F*_*s*_. The results are shown in Fig. 3b, where two notable features are discernible.

First, the trend of *τ*_*d*_ vs 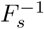 is visibly non-linear for the 3′end, the one with the widest range of 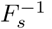. The non-linearity marks a further difference from the homopolymer case, where the simpler dissipative process yields 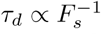, see above. Instead, the 3′ data are more compatible with two distinct linear regimes crossing over at *F*_*s*_ ~ 130 pN (inverse force of 0.76 · 10^*-*2^ pN^*-*1^).

Second, the *τ*_*d*_ data points are quite similar for the two ends. This is best appreciated in the inset of Fig. 3b which covers the region at small inverse forces (large *F*_*s*′_) for which data are available for both pulling directions. One notes that the 5 data have only small deviations from the low-force linear branch of the 3′-entry case.

The microscopic origin of both features are discussed further below.

#### Periodic driving

We next consider the xrRNA response to periodic driving. The setup is of interest for several reasons. First, it represents a still unexplored avenue where any novel insight can advance our understanding of RNA pore translocation. Second, it provides a term of reference for future electrokinetic experiments with e.g. solid state nanopores. Finally, the periodic driving mode is a simplified model for the action of enzymatic complexes that generally pull on the substrate in an intermittent and discontinuous manner[35–38]. These processive enzymes, which include exoribonucleases, are much larger than the xrRNA molecule of interest, which makes it computationally impractical to use them in place of the cylindrical pore.

We adopted a square-wave driving mode, with the pulling force switched between *F*_*p*_ (“on” phase) and 0 (“off” phase) at each semiperiod of duration *T/*2. For ease of comparison, we set *F*_*p*_ equal to 238 pN and 177 pN at the 5′ and 3′ ends, respectively, as these forces yield about the same *τ*_*a*_ ~4-5 10^3^ *τ*_MD_ at both ends in the static case. The period *T* was varied in the 0.0017-8.7 10^3^ *τ*_MD_ range, that is from being much smaller than *τ*_*a*_ to being comparable to it. Longer switching times were not considered because a significant fraction of translocations would otherwise complete already in the first “on” cycle.

Typical translocation curves at *T* = 5.2 · 10^3^ *τ*_MD_ are shown in Fig. 4. In this case, from two to three cycles are needed to activate the translocation process in half of the trajectories. The average activation times are equal to12.7 10^3^ *τ*_MD_ (5′-entry) and 8.6 10^3^ *τ*_MD_ (3′-entry), which are larger than the corresponding static values by more than a factor of 2.

**FIG 4.**
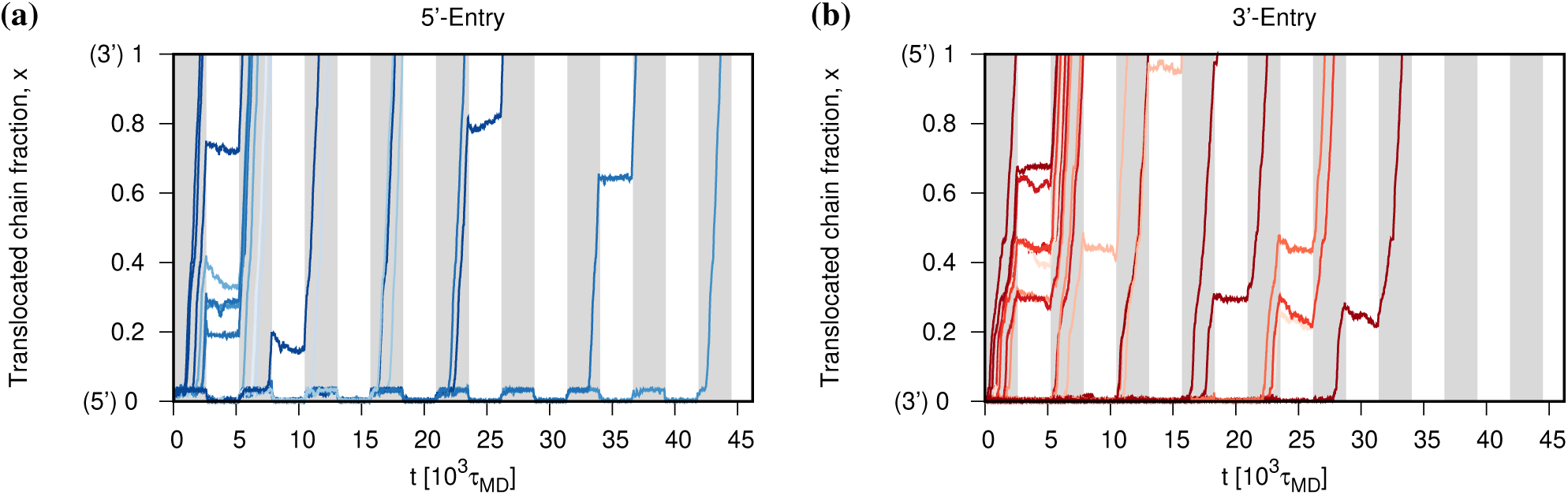
Translocation curves for periodic driving. Translocation curves for 5′ (blue) and 3^*′*^ (red) entries driven by a periodically switched force, *F* _*p*_. The latter was set equal to 238 pN and 177 pN at the 5′ and 3′ ends, respectively, so to have comparable activation times, see Fig. 3a. A shaded background highlights the semi-periods when the driving is “on”. In the “off” semi-periods, one notices pauses and even chain retractions from the pore.

Only few translocations complete in the same cycle where they initiate, and these cases are more common for 5′-entries. Most trajectories require two cycles to complete translocation after initiation. During the “off” phase of these trajectories, the translocation process is not only paused but can even regress. Note that most of the pauses and chain retractions from the pore occur in the first part of the translocation process, regardless of the pulling direction. This aspect will be revisited and discussed more in detail in the next section.

A systematic overview of how the periodic driving affects *τ*_*a*_ and *τ*_*d*_ is given in Fig. 5, where these quantities are plotted as a function of 1*/T*. In the static limit, corresponding to 1*/T* = 0, the activation times for 5′ and 3′ entries are about equal at the considered values of *F*_*p*_. As 1*/T* is increased from zero, *τ*_*a*_ increases too for both types of entries, though more prominently for the 5′ one, see Fig. 5a,c. A monotonic increase with 1*/T* is observed for the trajectory duration too, see Fig. 5b,d. In fact, *τ*_*d*_ approximately doubles going from the static case, 1*/T* = 0 to 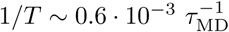 for both ends.

**FIG 5.**
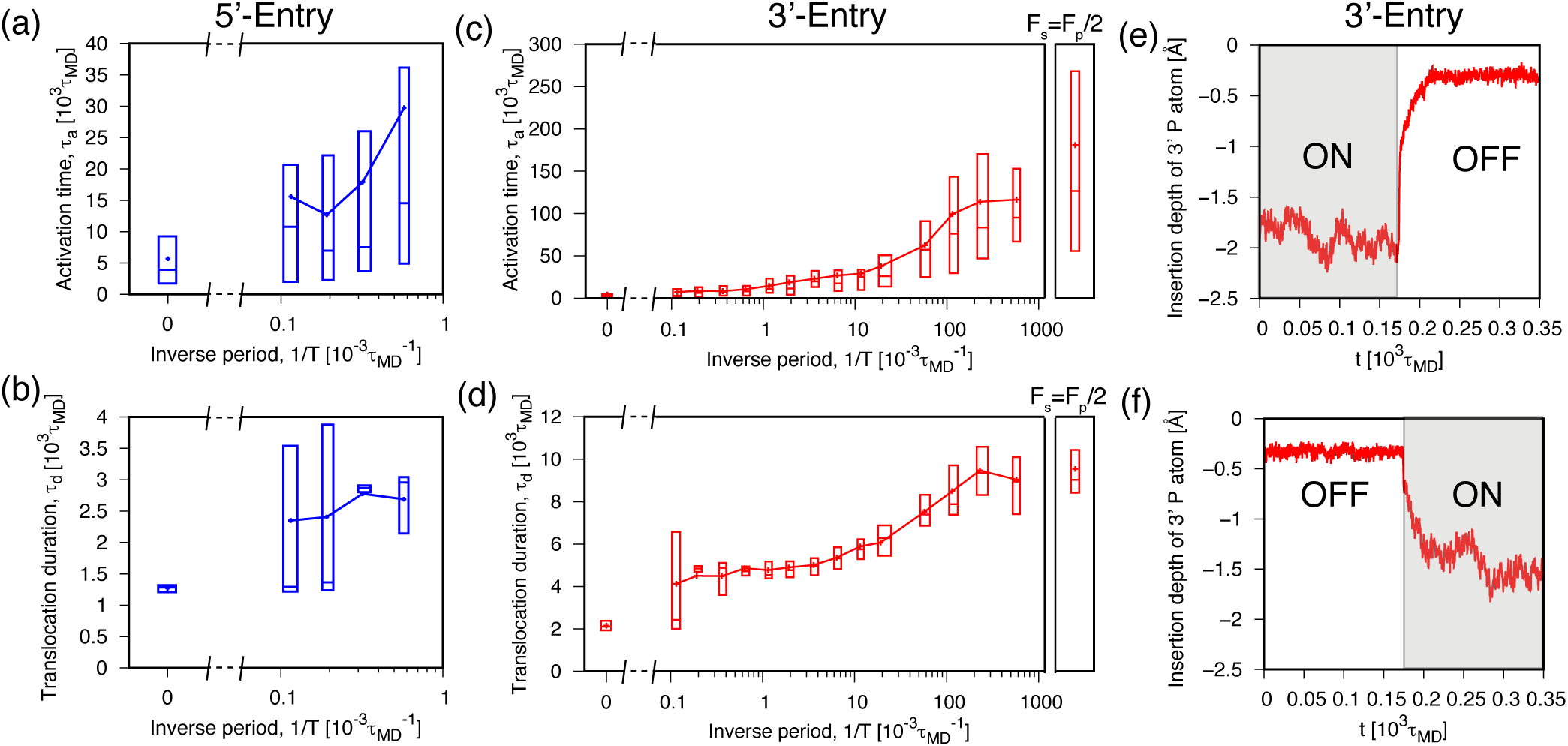
Translocation activation times and process duration for periodic driving. Box plots for (a) translocation activation times, *τ*_*a*_, and (b) translocation duration, *τ*_*d*_, for a periodically switched driving force, *F*_*p*_ = 238 pN, applied at the 5′ end. Corresponding plots for the 3′ end and *F*_*p*_ = 177 pN are given in panels (c) and (d). The box-plots at the right of these two panels show *τ*_*a*_ and *τ*_*d*_ for a static force equal to *F*_*s*_ = *F*_*p*_*/*2; the data are a reference for the asymptotic case 1*/T* → ∞, *T* → 0. The static case, instead, corresponds to 1*/T* = 0. Panels (e) and (f) show the system reponse to the sudden switching on or off of a static force *F*_*s*_ = 177 pN applied to the 3′ end of the xrRNA. The response is monitored through the pore insertion depth of the 3′ P atom. The box plot drawing convention is the same as for Fig. 3.

The spread of the distributions of *τ*_*d*_ and *τ*_*a*_ increases with 1*/T*, too. Both aspects reflects the occurrence of pauses and chain retractions during the intervening “off” phases which vary in number from one trajectory to the other and thus increase the duration and heterogeneity of the translocation process.

The limit 1*/T*→∞ is noteworthy because, when the switching is much faster than the characteristic response time of the system, one expects to recover the same behaviour as in the static case but with half the force, *F*_*s*_ = *F*_*p*_*/*2. We discuss this limit for 3′-entries only, for which the activation and duration times remain computationally addressable as the switching interval is reduced. The results are shown in Fig. 5c,d, where it is seen that, indeed, *τ*_*a*_ and *τ*_*d*_ become asymptotically compatible with the static values at half the force as *T*→0. Notice that the crossover towards the asymptotic limit occurs for *T* ~ 10 *τ*_MD_. This time duration is comparable to the system response time to a sudden switching of the pulling force *F*_*p*_, which is about 50 *τ*_MD_, see Fig. 5e,f. The results thus indicate that the half-force static response can be observed only for switching intervals smaller than the characteristic response time of the system.

#### Hindrance of secondary and tertiary elements

To locate specific xrRNA regions responsible for hindering translocation, we computed the so-called waiting time [29], *w*, of each nucleotide. The observable, which is experimentally relevant in connection with ionic current blockade, measures how long a nucleotide takes, on average, to cross the *cis* region and enter the pore, see Methods.

Typical waiting times profiles are shown in Fig. 6. The data are for different static forces applied at the two xrRNA ends, and are normalised by the average translocation duration, *τ*_*d*_, to facilitate comparison. The normalised *w* profiles differ significantly from 5′ entries and 3′ entries, but within each of these two sets, they are consistent across the considered forces, which yield significant variations of *τ*_*d*_.

**FIG 6.**
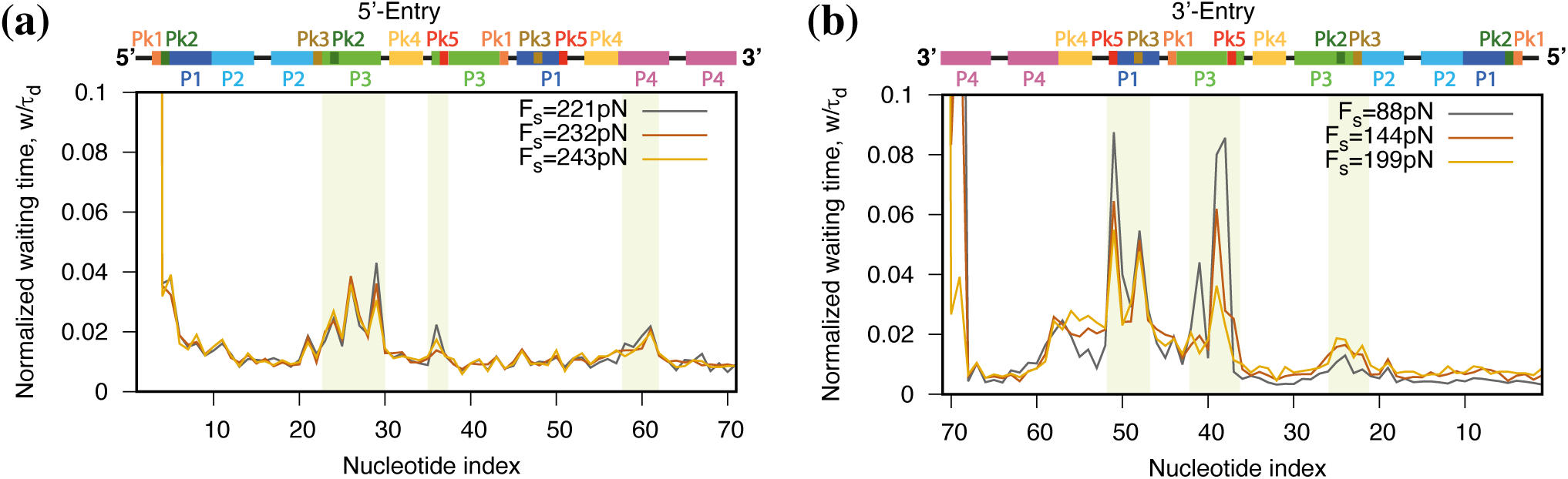
Site-dependent translocation hindrance; waiting times profiles. Waiting time profiles for three different static driving forces, *F*_*s*_, applied to the (a) 5′ and (b) 3′ ends. The waiting time, *w*, of a nucleotide corresponds to the average time required to cross the *cis* region and enter the pore. For ease of comparison, the *w* profiles are normalised to the average translocation duration. The off-scale values for nucleotides indexes smaller than 3 for the 5′-entry, and larger than 69 for the 3′-entry, reflect the typically large times required to activate translocation. The colored background highlights regions with the largest waiting times.

Qualitative differences for the two types of pore entries are not entirely unexpected, given earlier results on xrRNA directional resistance to translocation [8], which, however, hinged on the analysis of activation times. Instead, the present results highlight differences in waiting times, and thus provide a first insight on how diversely translocation proceeds from the two ends once it initiates.

We first discuss the *w* profiles for 3′ entries, Fig. 6b, which we could obtain for a wider range of forces thanks to the shorter translocation times. The largest resistance is offered by the stretch of nucleotides from U51 to A36 (ordered according to the 3′→ 5′ translocation direction), where two sets of peaks are observed. The first one involves Pk3 and Pk5 and the second involves the 3′ arms of helix P3. The height of these peaks is highest at low force. The remainder of the xrRNA structure beyond A36 offers relatively little hindrance, except for the neighborhood of Pk2.

The *w* profiles for 5′ entries, which take much longer to translocate, were, by computational necessity, collected at higher forces, see Fig. 6a. The first encountered, and most prominent peak corresponds to the 5′ arm of helix P3, which includes the neighborhoods of Pk3. Two further minor peaks are observed close to the 5′ arm of Pk5 and helix P4.

The very different nature of the profiles at the two ends can be rationalised by considering the sequential order in which secondary elements are disrupted by translocation, and especially that helices can offer resistance only for the first translocating arm, after which they become fully unzipped.

These observations suffice to account for the qualitative features of the *w* profiles for both types of pore entries. For 3′ entries, translocation initiates with the unzipping of the P4 helix. After this event, which is off-scale in Fig. 6b, the first appreciable hindrance is encountered in correspondence of Pk3-Pk5. Translocation next proceeds with the disruption of the contacts in helix P3. Once the latter is unzipped, the remainder xrRNA structure is mostly void of secondary elements, the residual ones being only Pk2 and P2 that translocate with little resistance. From a quantitative point of view, it is interesting that, although the applied static force is constant throughout the translocation process, all helices P1 to P3 are disrupted in a small fraction of the time required to activate the unzipping of helix P4.

Analogous considerations apply to the 5′ end, Fig. 6a. Here translocation initiates with the disruption of Pk1 and Pk2 (again, this event is off scale in the graph of Fig. 6a). No particular hindrance is found for P1 and P2, while more resistance is offered by the 5 arm of P3. After P3 becomes unzipped, the only significant remaining secondary element is helix P4 and, in fact, translocation proceeds unhindered up to this point, which defines the last obstacle of the process.

The single feature of the *w* profiles that bears a noticeable force dependence is the height of the peaks in the 3′ arms of helices P1 and P3 of Fig. 6b (3′ entry). The height of the peaks is strongly diminished when the applied force is increased. It is physically appealing to associate the decrease of normalised waiting times with a reduction of the free-energy barriers hindering translocation. We accordingly surmise that the crossover between the two different linear regimes for *τ*_*d*_ in Fig. 3b follows from a change of the unzipping barriers for P1 and P3. In support of this speculation we provide two observations. First, for *F*_*s*_ ~144 pN the peaks have an intermediate height between the maximum and minimum values, and this force is comparable to where the *τ*_*d*_ crossover is observed, for *F*_*s*_ ~130 pN. Second, the milder slope of the high-force (small inverse forces) branch of *τ*_*d*_ is consistent with translocation barriers becoming smaller, as the peak waiting times, at large *F*_*s*_.

## SUMMARY AND CONCLUSIONS

Nanopore translocation is a powerful single-molecule probing technique that has been used in diverse contexts: from studying the physical response of homopolymers[19–21, 32, 34, 39], to the topological friction in chains with knots and links[33, 40–48], for sequencing and analyzing biopolymers’ secondary and tertiary structures [7, 11, 29, 41, 49–57], and study RNA too[1–4, 6–8].

Here, we used nanopore translocation simulations to study the compliance of a viral RNA to be driven through a narrow pore. We focused on the 71-nucleotide long xrRNA from the Zika virus. In a previous atomistic study from our group[8], the translocation response of the xrRNA was studied with a force-ramping protocol, a common setup for force spectroscopy experiments. The xrRNA was found to be capable of withstanding much higher pulling forces at the 5′ end than at the 3′ one before translocation could initiate. The strongly directional resistance was ascribed to the particular architecture of the xrRNA, which tightens, thus offering more resistance, when its 5′ region is pulled against the pore surface. The observed directional resistance is arguably harnessed by the molecule to elude the degrading action of cellular exonucleases, which engage RNAs from the 5′ end[13–15].

Here, we used atomistic simulations and a native-centric model [24, 25] to clarify three different aspects of RNA pore translocation: how the onset and duration of RNA translocation depend on the driving force, what are differences of using static or periodic driving modes, and how the progress of translocation is affected by secondary and tertiary elements.

We established the following results. First, the start of xrRNA translocation is so delayed with respect to the time of application of the driving forces, that translocation activation times can exceed by orders of magnitude the duration of translocation process itself. In the static case, the force-dependent duration of the delays is compatible with a two-state activated process involving barriers of very different widths at the two ends. The barrier widths are not dissimilar from those estimated previously with the Bell-Evans analysis of force-ramped trajectories[8] and the widest barrier (5.6 ±0.2 Å) is encountered at the 5′ end. This implies that the relative difference in activation times at the two ends grows exponentially as the pulling force is lowered.

Second, we observe that using a periodic driving instead of a static one increases significantly both the activation times and the duration of the process. The variance of both quantities are increased with respect to the static case too. Both aspects are accounted for by the fact that the process is stalled and can even regress during the off phase of the driving cycle. We thus conclude that the directional resistance of xrRNA is enhanced by discontinuous pulling modes, such as those that arguably occur *in vivo* when the xrRNA is engaged by processive enzymes, such as exonucleases, replicases and reverse-transcriptases.

Finally, we investigated the non-uniform progress of translocation once started. Regardless of the pulling directionality, the largest hindrance is always encountered in the first part of the trajectory. Analysis of the waiting times profiles provides a simple rationale for this result: secondary elements such as helices can offer resistance only for one of their two arms, the one pulled first into the pore, after which they become fully unzipped. Consequently, as translocation progresses, the RNA becomes rapidly depleted of intact secondary elements and the process can proceeds with less and less hindrance. More quantitatively, we pinpointed the specific xrRNA regions most responsible for hindering translocation. These regions are different for the two pulling ends but, in both cases, they involve helix P3 which includes pseudknotted nucleotides.

The findings are of interest and have implications beyond the case of Zika xrRNA, as they highlight the several ways in which the translocation of folded RNAs differs from that of homopolymers. For general models of homopolymers, which are exclusively informed by chain connectivity and excluded volume interactions, translocation can initiate concomitantly with the application of the driving force and then proceeds smoothly following an asymptotic scaling law defined by the metric exponent[32, 34]. We instead observe that folded RNAs, which are stabilised by specific intramolecular interactions, present major delays in the start of translocation. In fact, the activation times can exceed by far the duration of translocation itself. In addition, rather than proceeding smoothly, xrRNA translocations present slowing downs, pauses and even retractions that, though depending on pulling directionality, occur in correspondence of specific secondary and tertiary elements.

The results complement and generalize previous studies of translocating RNA hairpins[52, 58, 59] and offer valuable terms of reference for future theoretical and experimental studies. In particular, the results ought to be useful to validate simpler RNA models, e.g. based on coarser structural descriptions and effective intramolecular interactions, which would be more amenable to numerical characterization and hence more widely applicable. Natural targets of such endeavours would be other viral RNAs, especially those already known to be capable of resisting degradation by exonucleases [14, 60, 61]. In addition, we expect that the present elucidation of the interplay of translocation directionality, pulling mode, and RNAs’ secondary and tertiary structures, ought to be useful for interpreting and, possibly, designing future single-molecule experiments. For both theory and experiment, we believe that a promising avenue would be to extend considerations to how exactly processive enzymes engage and translocate RNAs with complex architecture.

## ACKNOWLEDGEMENTS

This work was partially supported by MUR, the Italian Ministry of University and Research. We acknowledge the CINECA award under the ISCRA initiative, for the availability of high performance computing resources and support (“UDNANO” project, code HP10CE4Q04).

